# Perceptual Novelty in Tinnitus a Causative Factor for its Persistence. A Stimulus Novelty Based P300 Paradigm on Acute, Chronic, and Non-Tinnitus Controls

**DOI:** 10.1101/2025.08.15.670598

**Authors:** Abishek Umashankar, Kai Alter, Phillip E. Gander, William Sedley

## Abstract

Our understanding of tinnitus pathophysiology may be greatly advanced by understanding how the condition evolves from its initial onset or acute stage to its chronic manifestation. Such a transition likely reflects dynamic neurophysiological changes within central auditory and non-auditory networks. Previous studies have highlighted that individuals with acute tinnitus tend to have increased activity in the regions of anterior cingulate cortex, inferior parietal lobe, and insula all of which are essential in constituting the salience network. We therefore aimed at tapping into the salience network of tinnitus through a novelty based P300 paradigm in individuals with Acute, Post Acute (six months follow up since tinnitus onset) Chronic, and Controls. Participants were presented with an auditory oddball paradigm comprising three deviant types: (1) *novel* environmental sounds, (2) *low-frequency* tonal deviants, and (3) *high-frequency* tonal deviants, embedded within a sequence of frequent standard tones. Our results indicate a significant drop in P300 amplitude during the Post Acute stage across the three deviant stimuli, highlighting the substantial influence of the anterior cingulate cortex/salience network in possible generation of tinnitus and inferior parietal lobe in the persistence of tinnitus.

## Introduction

Tinnitus, an auditory phantom perception, has a remission rate of just 15%, which is considerably lower than that of other disorders (Gopinath et al., 2010). This indicates that for most individuals acquiring Tinnitus, there is an early transition from the onset of tinnitus to its chronic phase, which generally persists over time. Additional research indicates that tinnitus lasting over four weeks, in the absence of Acute external or middle ear diseases, is likely to continue long-term in at least 80% of persons (Vielsmeier et al., 2020). However, the question remains as to why tinnitus predominantly remains chronic and does not resolve, despite the brain’s intricate systems designed to achieve homeostasis via different gain control mechanisms. In earlier our previous research, we thoroughly examined the alterations in central gain from the onset to the chronic stages of tinnitus. We found that, although reductions in hypersensitivity are observed through Intensity Dependence of Auditory Evoked Potentials (IDAEP) (Umashankar, Sedley, et al., 2025), the gain associated with subjective sound sensitivity remains unchanged over time (Umashankar, Gander, et al., 2025).This implies that the mechanisms contributing to the chronicity of tinnitus may commence early due to no variations seen between the onset and chronic stages of tinnitus.

This was supported by the opinions of both Hullfish et al. (2019) and Noreña & Farley (2013) where both highlight that changes related to the early chronification of tinnitus begin soon after the onset of tinnitus with even Hullfish et al. (2019) further highlighting that acute stages of tinnitus may be less than 4 weeks. It is also further to note that the processes that initiate tinnitus are not necessarily the same as those that make it chronic as tinnitus may be initiated by temporary events like noise exposure, but its persistence involves complex changes in the brain’s auditory pathways, leading to a chronic state (Kaltenbach, 2007; Kaltenbach & Manz, 2012; Sedley et al., 2016). It is further to note that the homeostatic processes in the brain may mask tinnitus-related changes over time as highlighted by Yang et al. (2011) and Roberts et al. (2013) where they further highlight that there are mechanisms of adaptation following the onset of tinnitus which can be evidently seen why tinnitus severity does not always correlate with hearing loss. In addition to alterations in homeostatic processes from the onset of tinnitus to its chronic state, numerous changes associated with chronic tinnitus may result from the condition itself rather than the causal mechanisms as discussed by Umashankar, Gander, et al. (2025) where increased distress at the time of tinnitus onset may contribute to its chronification, primarily relating to the reactivity to tinnitus rather than its underlying cause.

In summary, further exploration is warranted of potential mechanisms contributing to the persistence of tinnitus, which might involve wider networks than just the auditory pathway. Investigating alterations in brain activity along the development of tinnitus and its subsequent persistence is crucial to understanding the low remission rate of tinnitus. A limited number of source power or functional connectivity studies have compared acute and chronic tinnitus, using resting state EEG/fMRI measurements. Lan et al. (2021) identified a notable enhancement in power within the middle frontal gyrus and parietal cortex in the gamma band for acute tinnitus in comparison to the chronic tinnitus group. Cai et al. (2020) conducted a functional connectivity study that demonstrated an enhancement in both functional and causal connectivity for the superior temporal gyrus, amygdala, and nucleus accumbens in acute tinnitus group relative to persons without tinnitus. Both these studies compared separate acute and chronic tinnitus groups cross-sectionally. Extrapolating from research findings not directly examining acute tinnitus, Vanneste et al. (2011) proposed that patients with newly developed tinnitus exhibit heightened activity in the auditory cortex, supplementary motor area, dorsal anterior cingulate cortex, and insula. De Ridder et al. (2022) proposed a triple network model for tinnitus, suggesting that persons with Acute Tinnitus exhibit an anti-correlation between the default mode network and the central executive network, which facilitates the propagation of tinnitus distress through the salience network. To summarise the limited findings, electrophysiological and imaging results indicate an increase in activity in non-auditory regions around the start of tinnitus compared to its chronic stages. This may manifest as, or result from, heightened distress or increased focus of attention on the tinnitus at its onset. Much current tinnitus theory emphasises the involvement of the pregenual or dorsal/lateral anterior cingulate cortex in the onset of tinnitus, either by activating tinnitus or by exhibiting heightened activity associated with newly developed tinnitus characterised by distress (Hullfish et al., 2019; Song et al., 2015; Vanneste et al., 2024). Other literature emphasises the significance of insular activity during acute phases of tinnitus. Chen et al. (2023) demonstrated atrophy of the ventral anterior insula in chronic tinnitus compared to acute tinnitus groups and concluded that the generation and persistence of tinnitus result from the aberrant structural and functional reorganisation of the insula. Araneda et al. (2018) emphasised the significance of the prefrontal cortex in the onset and maintenance of tinnitus, attributing to it deficiencies in the executive functions and inadequate inhibitory regulation. Furthermore, acute Tinnitus literature have highlighted in common increased oscillatory activity to be around regions like the anterior cingulate, insula, and prefrontal cortex. All of these regions together are essential in constituting the salience network.

Both salience (prominence/significance) and novelty (newness or unexpectedness) might be strongly associated with acute tinnitus due to prominence and newness the tinnitus has during its onset. In the present study, therefore, we sought to assess the role of the salience network, to potentially explain the onset of tinnitus. However, another question pertains to whether the factors of salience or novelty processing contribute to the persistence and non-remission of tinnitus. Hence, to commence an investigation into the mechanisms underlying the persistence of tinnitus, it would be prudent to examine the involvement of the salience network in relation to the salience or novelty of tinnitus. One way would be to assess novelty mechanisms of tinnitus through an electrophysiological P300 paradigm, where the spatiotemporal dynamics are centred around the anterior cingulate and the temporoparietal networks (Yago et al., 2003). The P3a, or novelty-driven P300, along with the P3b, engages the ventral frontal/parietal attention network respectively, with the former designed to detect salient changes and the latter focused on attention orientation, both elicited by the ACC and temporo-parietal regions (Kim, 2014). Based on the aforementioned literature reporting increased activity of non-auditory regions in the prefrontal cortex regions during acute stages of tinnitus, we hypothesize that the components of salience or novelty would be heightened during the acute phases, subsequently diminishing and stabilising throughout the ensuing chronic stage and this follows with the other components of P3b too. We further hypothesize that the enhanced salience during the onset of tinnitus may further result in activating networks that might result in the persistence of tinnitus as few studies highlight the role of attentional networks towards the tinnitus to contribute to the persistence of tinnitus (Haider et al., 2018).

## Methods

### Participants

A total of 23 participants with Acute Tinnitus (symptoms lasting for less than 4 to 6 weeks) were subsequently followed up 6 months later as they transitioned into the Chronic stage. For convenience, this group is referred to as the Post Acute Tinnitus group to distinguish it from the actual Chronic Tinnitus data collected cross-sectionally in this study, 27 participants with Chronic Tinnitus (symptoms persisting for more than 6 months), and 19 non-Tinnitus participants (no Tinnitus symptoms) participated in the current study, recruited as a community sample through advertisements on Google adverts and internally at Newcastle University. Individuals aged 18 and older capable of providing informed consent were included, but those with Meniere’s disease, epilepsy, or middle ear pain or infections were excluded from the study.

Control participants were matched individually to those in the Acute Tinnitus group for age, sex, and hearing. Control participants were presented with stimuli that matched the frequency and presentation ear(s) of their matched tinnitus participants. An attempt to match the Chronic Tinnitus group to the other groups in these regards was only partially successful.

Details are displayed in Table 1.

**Table 6.1:** Participants’ demographics for each group. N denotes number of samples, M denotes mean, SD denotes Standard Deviation, MD denotes Median, F statistic indicates a One Way ANOVA has been carried out, X^2^ indicates a chi-square goodness of fit test, z test indicates non parametric Man-Whitney U test, NS denotes No Significance. * indicates presence of statistical significance at p < 0.05. Hearing thresholds were calculated by averaging the left and right ear values.

| Parameters | Acute | Chronic | Controls | Significance | Post hoc |
| --- | --- | --- | --- | --- | --- |

|  | **Tinnitus** | **Tinnitus** |  | **Overall** | **Significance** |
| --- | --- | --- | --- | --- | --- |
| **Age (Years)** | N = 22<br>M = 55.36<br>SD = 13.65 | N = 24<br>M = 60.54<br>SD = 8.63 | N = 19<br>M = 56.26<br>SD = 9.23 | F = 0.228<br>p = 1.514 | NS |
| **Gender (M/F)** | 15/7 | 18/6 | 11/8 | $X^2 = 1.422$<br>p = 0.491 | NS |
| **Hearing (1 kHz) dBHL** | M = 11.93<br>SD = 11.15 | M = 13.33<br>SD = 7.89 | M = 10.39<br>SD = 6.47 | F = 0.593<br>p = 0.556 | NS |
| **Hearing (4 kHz) dBHL** | M = 25.68<br>SD = 16.33 | M = 38.75<br>SD = 14.52 | M = 21.32<br>SD = 15.03 | F = 7.776<br>p < 0.001* | Acute –<br>Chronic; p = 0.014*<br>Chronic –<br>Control; p = 0.001* |
| **Hearing (8 kHz) dBHL** | M = 39.09<br>SD = 19.67 | M = 49.79<br>SD = 19.22 | M = 27.5<br>SD = 18.54 | F = 7.173<br>p = 0.001* | Chronic =<br>Control; p = 0.001* |
| **Tinnitus Pitch (Hz)** | M = 6823.65<br>SD = 1459.67 | M = 6865.27<br>SD = 1989.12 | M = 6476.5<br>SD = 1206.27 | F = 0.354<br>p = 0.703 | NS |
| **Tinnitus Loudness (dBSL)** | M = 11.46<br>MD – 8.33<br>SD = 12.33 | M = 5.34<br>MD – 4.5<br>SD = 10.59 | N/A | z = 1.617<br>p = 0.106 | N/A |

### Audiological Assessment

Prior to the experiment, participants completed a pre-screening form containing questions designed to confirm eligibility based on the inclusion and exclusion criteria. The experimenter, a qualified clinical audiologist with experience in tinnitus, conducted interviews with the participants prior to the experiment, confirming a medical history consistent with subjective tinnitus and the lack of atypical symptoms or concerns about an unidentified underlying cause. Participants provided demographic information, including age and sex, as well as information about their tinnitus, such as type (tonal, noise-like, or other), duration, which ear(s), and a past physical and mental health history, including any otological disorders. Four common, validated questionnaires—the Tinnitus Handicap Inventory (THI) (Newman et al., 1996), the Tinnitus Functional Index (TFI) (Meikle et al., 2012), the Hyperacusis Questionnaire (HQ) (Khalfa et al., 2002), and the Inventory of Hyperacusis Symptoms (IHS) —were used to evaluate the impact and distress related to tinnitus and any co-existing hyperacusis symptoms (Greenberg & Carlos, 2018).

Participants underwent a Pure Tone Audiometry (PTA) test to establish hearing thresholds at octave frequencies ranging between 250 Hz up to 8 kHz. Thresholds were estimated based on the initial presented level at either 40 dB HL or above depending on the participant’s residual hearing. If perceived audible, the stimulus was continuously reduced by 15 dB HL until the participant did not perceive it as audible. From the first reversal, a 5 dB up, 10 dB down staircase procedure was then used until the final threshold was established by two positive responses out of three trials (Carhart & Jerger, 1959).

### Tinnitometry

Tinnitus pitch and loudness were determined via a *tinnitometry* procedure, for participants with either acute or chronic tinnitus. In cases of unilateral tinnitus, matching stimuli were presented contralaterally, and in bilateral tinnitus they were presented bilaterally. Participants first matched their tinnitus for loudness and then for frequency from a starting reference stimulus of 6 kHz. In bilateral cases where the tinnitus was asymmetric, additional adjustments were made to balance the two ears. Both loudness, then frequency, were iteratively adjusted until the sound closely matched the participant’s tinnitus, and no further changes were judged necessary by the participant. This procedure was repeated across three trials and a mean of the trials was taken and considered as the final loudness and pitch of the tinnitus. The stimulus used for this procedure was either a pure tone or narrow band noise (1/3 octave with Hanning spectrum) depending on the subjective similar bandwidth of the tinnitus.

### P300 Stimulus

Four categories of stimuli based on different acoustical frequencies were delivered for the P300 study. The High Frequency Deviance (HFD) which is a pure tone stimulus presented at the participant’s tinnitus frequency, Standard Stimuli (SS) which is a pure tone stimulus presented at one octave below the tinnitus frequency, Low Frequency Deviance (LFD) which is a pure tone stimulus presented at one octave below the standard stimuli, and the Novel Stimuli (NS) which is comprised of a collection of random ambient sounds (e.g., dog bark, cat meow, car honk), with each sound given singularly to maintain novelty, and all sounds were specifically chosen to ensure neutral valence (emotional neutrality in response to the stimulus such as white noise, bird chirps, water droplets, etc). All four stimuli were presented at a 60% level relative to the differences between the Uncomfortable Loudness Level (ULL) and the threshold Dynamic Range (DR) with NS priorly undergoing an RMS match to the average of the first three categories before the ULL and DR were established.

Both the threshold and ULL were determined via an ascending procedure with the threshold established at 50% response based on the number of trials and the ULL established independently to prevent the presentation of multiple sounds at potentially uncomfortable elevated intensities. For the controls, an individual was matched to the Acute Tinnitus group where the respective frequency of that person was taken as HFD, SS, and LFD. The dynamic range of those frequencies was later calculated for the controls in the same way as for Tinnitus participants.

### P300 Stimulus Parameters and Recording Parameters

EEG was recorded using a 64-channel Active Two Biosemi system in a soundproof room. These 64 channels were sufficient to remove ocular artifacts and an additional ocular channel was not used. Electrode offset was kept at the manufacturer’s recommended limits of ±10 mv. A total of 1200 standards and 50 deviants of each type were presented in a randomized order with 5 to 11 standards between deviants. The HFD, SS, NS, and LFD all had a 10 msec Onset/Offset Ramp and 300 msec duration of presentation and across the four Stimulus, an 800 msec Interstimulus Interval was maintained. With respect to the intensity presentation of NS, the intensity of the novel stimulus was calibrated to correspond with the average RMS (Root Mean Square) across all the NS based on a sampled loudness matched stimulus. By standardising the RMS, all stimuli were delivered at a uniform loudness level, facilitating a clear comparison of the P300 response to novelty devoid of intensity-related fluctuations. The experiment was a Go-No-Go paradigm where the participants were asked to positively respond to the HFD and LFD stimuli only, by pressing the spacebar button every time they heard either of these, and to ignore the SS and NS stimuli. A fixation cross (+) that changes into a X symbol every time they pressed the spacebar was provided as visual feedback for their response. All stimuli were generated and presented in Matrix Laboratory (MATLAB) version R2019a. The total duration of the experiment was approximately 25 minutes.

### EEG Pre Processing

All EEG data underwent preprocessing in MATLAB version R2019a EEGlab toolbox. Firstly, re-referencing at P9/P10 (Linked mastoids) was carried out and later filtering was performed with a high pass of 1 Hz and low pass of 30 Hz. This was followed by epoching between - 0.1 and 1 s and channel rejection based on visual inspection and keeping 0.8 as the accepted correlation between channels. Artifact rejection was done using the Independent Component Analysis (ICA) where artifact-containing components like an eye blink, eye movements, muscle artifacts, and other electrical artifacts, were removed. Apart from ICA, artifact rejection was also carried out using individual trial rejection where components are auto rejected keeping a probability of 5 and Kurtosis of 8 as the threshold. Baseline correction was further done from −100 to 0 ms from stimulus onset.

### Data analysis

The P300 component was produced and evaluated from the respective epochs for the Acute, Post Acute, Chronic Tinnitus, and Control groups. The total number of trials was averaged for each condition, and the P300 response was measured at a post-stimulus interval of a time window between 250 ms to 400 ms for LFD and HFD, and between 220 ms to 400 ms for NS, specifically at the Pz electrode for LFD and HFD, and at FCz for NS.

Automatic peak height detection was conducted for the aforementioned time windows corresponding to the respective P300 groups. A visual inspection of each waveform under every situation was conducted to identify a suitable P300 response and subjects were rejected if the P300 peak/waveform was not visually present at the above-mentioned latency windows for each condition.

### Statistical Analysis

Statistical analysis was performed using the Statistical Package for Social Science (SPSS) with descriptives and inferential statistics calculated. Upon the Shapiro-Wilk test of normality, the data was normally distributed across the three groups, and hence parametric statistics were used. We used P300 relative peak amplitudes for our statistical analysis.

Relative peak amplitude was defined by calculating the average amplitude across three conditions and then normalizing each condition by dividing its amplitude by this average to obtain the relative amplitude for each condition. The rationale for this is the presence of subject-specific variations in amplitude across all conditions. Examining the relative differences in salience and attentional processing of novel, non-tinnitus-related, and tinnitus-related stimuli aids in mitigating these absolute differences. For the within-group Acute-Post Acute comparison, we employed a method to establish the relative amplitude by calculating the average amplitude for all three conditions in each subject for both time points combined and then normalizing each condition by dividing its amplitude by this average. This was to give a more equal weighting to each subject, considering the large between-subject differences in overall P300 amplitude.

We conducted a One-Way ANOVA with subject groups as the independent variable and amplitudes across conditions (HFD, LFD, and NS) as the dependent variable to determine the main effects of the groups and to analyse within-group differences; a Tukey’s post hoc test was subsequently performed. A paired t test was conducted to determine the significant difference between the paired groups of Acute and Post-Acute Tinnitus.

### Source reconstruction

Source reconstruction was performed across groups and conditions for the P300 amplitude. Relative amplitudes were employed in the statistical analysis, necessitating normalisation for the source analysis as well. As a part of normalization, for each subject, the trial-averaged time series for every electrode was divided by the mean P300 amplitude value across conditions and subjects within that group for the FCz electrode between time point 220ms and 400 ms for NS and Pz electrode between 250 ms and 400 ms for both LFD and HFD. Source reconstruction was conducted initially only on group comparisons that demonstrated statistically significant difference in sensor space (differences in relative amplitude), i.e. source reconstruction was for localisation purposes, of between-group differences already found to be significant.

To compare source-localized P300 activity among Acute Tinnitus, Chronic Tinnitus, and Control groups, we first identified statistically significant differences at the sensor level—i.e., the scalp-recorded event related P300 signals. Upon detecting such significance, we employed standardized low-resolution brain electromagnetic tomography (sLORETA) to perform voxel-by-voxel independent t-tests, enabling the localization of neural generators underlying the observed P300 components. The analysis was done sing SLORETA’s built-in statistical tools individually between the groups (Pascual-Marqui et al., 1994). These tests identified regions where cortical activation differed significantly between groups.

Additionally, to the targeted source analyses based on significant sensor-level differences, paired t tests were conducted for within-group comparisons across different stimulus conditions (LFD, HFD, and novelty) to assess condition-specific differences in P300 source activation in case there was a paired differences at source space (not evident in sensor space). Non-parametric randomization tests (Nichols & Holmes, 2002), using 5000 permutations, were applied to control for multiple comparisons across voxels. Statistical maps were established regardless of significant difference (as significant differences have already been established at sensor space) and significant voxels were grouped based on their corresponding brain regions, including cortical lobes and Brodmann areas. It is further to note that maximum t statistics levels within the time series during group comparisons were considered as the region of cortical activation for that group comparison.

## Results

Out of the total participants enrolled (Acute Tinnitus – 23, Chronic Tinnitus – 27, Controls – 19), five individuals were omitted owing to data quality: one from the Acute group, three from the Chronic group, and one from the Post Acute group. The final participant composition consisted of 22 individuals with Acute Tinnitus (mean tinnitus duration: 4.27 weeks, median: 4 weeks, SD: 2.45 weeks), of which 4 participants exhibited a tinnitus duration ranging from 7 to 10 weeks and were classified as outliers. Among the 22 individuals with Acute Tinnitus, 9 returned after 6 months to form the Post Acute Tinnitus group, 24 with Chronic Tinnitus (mean tinnitus duration: 8.73 years, median – 4 years, SD – 9.14 years) and 19 Controls.

### Demographics and Hearing

Efforts were made to match participant groups based on age, sex, and hearing ability. Participants were matched for age (F(2,62) = 1.514, p = 0.228), sex (*X^2^* = 1.422, p = 0.491) and with respect to hearing, no significant main effect was observed for hearing thresholds at 1 kHz (F(2,62) = 0.593, p = 0.556). A significant main effect was present at 4 kHz (F(2,62) = 7.778, p < 0.001), with a notable difference between Acute Tinnitus and Chronic Tinnitus (p = 0.019) and between Chronic Tinnitus and Controls (p = 0.001). Chronic Tinnitus exhibited higher thresholds (M = 38.75, SD = 14.52) compared to Acute Tinnitus (M = 25.68, SD = 16.33) and Controls (M = 21.31, SD = 15.03). No differences were noted between Acute Tinnitus and Controls (p = 0.64) Comparable outcomes were observed at 8 kHz, revealing a significant main effect (F(2,62) = 7.173, p = 0.001), with notable differences solely between Chronic Tinnitus and Controls (p = 0.001). Chronic Tinnitus exhibited a higher threshold (M = 49.79, SD = 19.22) compared to Controls (M = 27.5, SD = 18.54). No significant changes were noted between Acute Tinnitus and Chronic Tinnitus (p = 0.169), nor between Acute Tinnitus and Controls (p = 0.126). Figure 1 illustrates the hearing thresholds, while table 1 provides information regarding the participants’ demographics.

**Figure 1.**
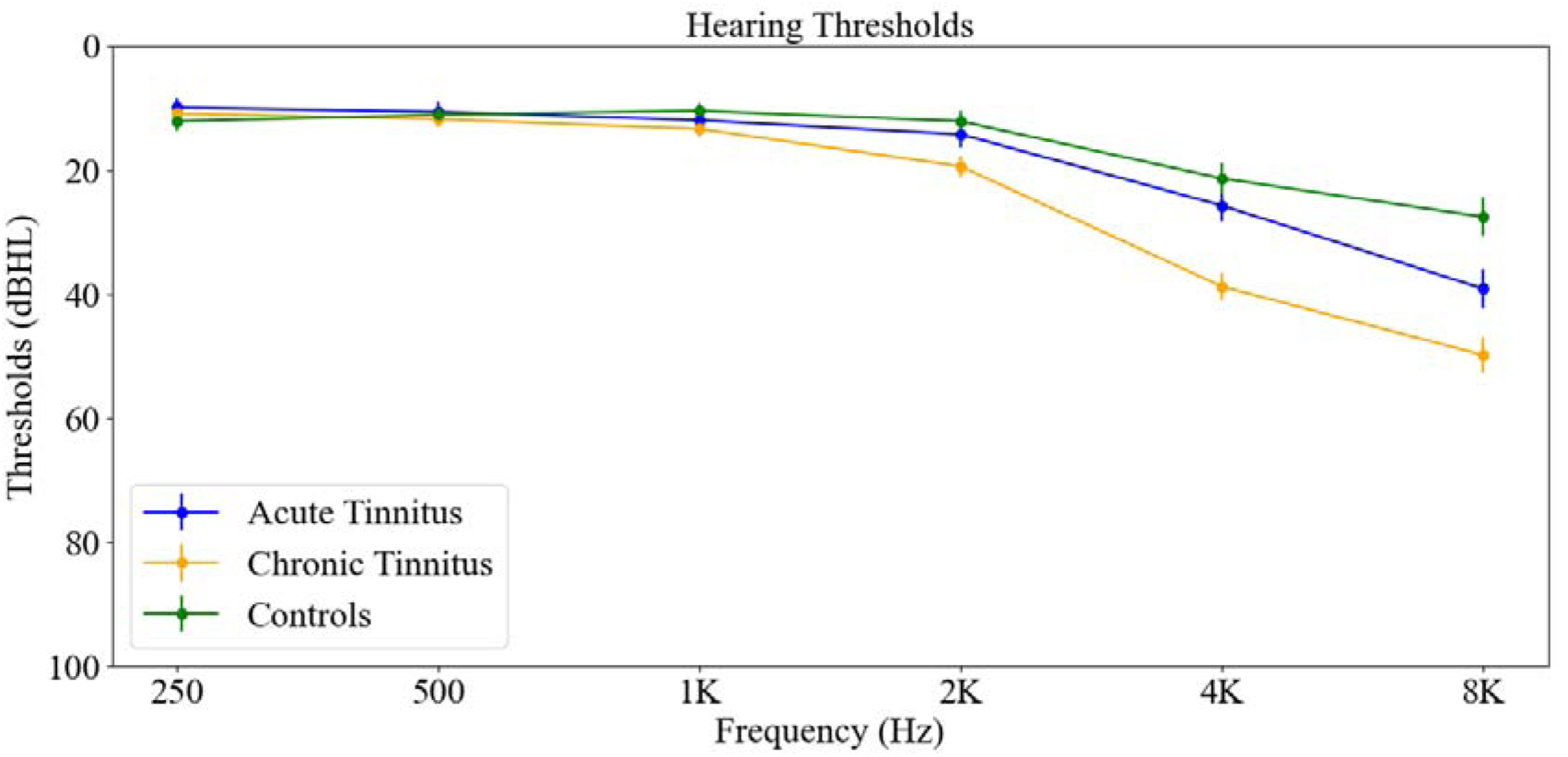
Combined average hearing threshold for both ears across Acute, Chronic, and Non-Tinnitus Controls. Acute Tinnitus and Controls were matched for hearing at 1 kHz, 4 kHz, and 8 kHz. However, Chronic Tinnitus was not matched for 4 kHz and 8 kHz (refer table 6.1).

### Symptom scores

Symptom scores for tinnitus and hyperacusis were obtained and calculated cross sectionally across the groups. With respect to the tinnitus symptom scores, no significant differences were obtained between the Acute and Chronic Tinnitus for THI (t(40) = −1.254, p = 0.217) and for TFI (t(40) = −1.091, p = 0.282). Regarding the hyperacusis questionnaires, there were no significant differences observed for the HQ questionnaire scores across the Acute, Chronic, and Control groups. (F(2,57) = 2.877, p = 0.064. However, a significant effect was obtained for IHS (F(2,57) = 7.177, p = 0.001) across the groups with a significant difference between Chronic Tinnitus and Controls (p = 0.001) with Chronic Tinnitus (M = 50.45, SD = 17.35) having increased hyperacusis scores when compared to controls (M = 33.11, SD = 8.27). No differences were obtained between Acute Tinnitus and Chronic Tinnitus (p = 0.188) and between Acute Tinnitus and Controls (p = 0.116).

For the longitudinal changes, there were no significant differences between the Acute and Post Acute Tinnitus for THI (z = −1.5, p = 0.136), TFI (z = −1.23, p = 0.219), HQ (t(5) = - 0.169, p = 0.873), and IHS (z = −0.315, p = 0.753). Refer table 2 for further details.

**Table 2:**
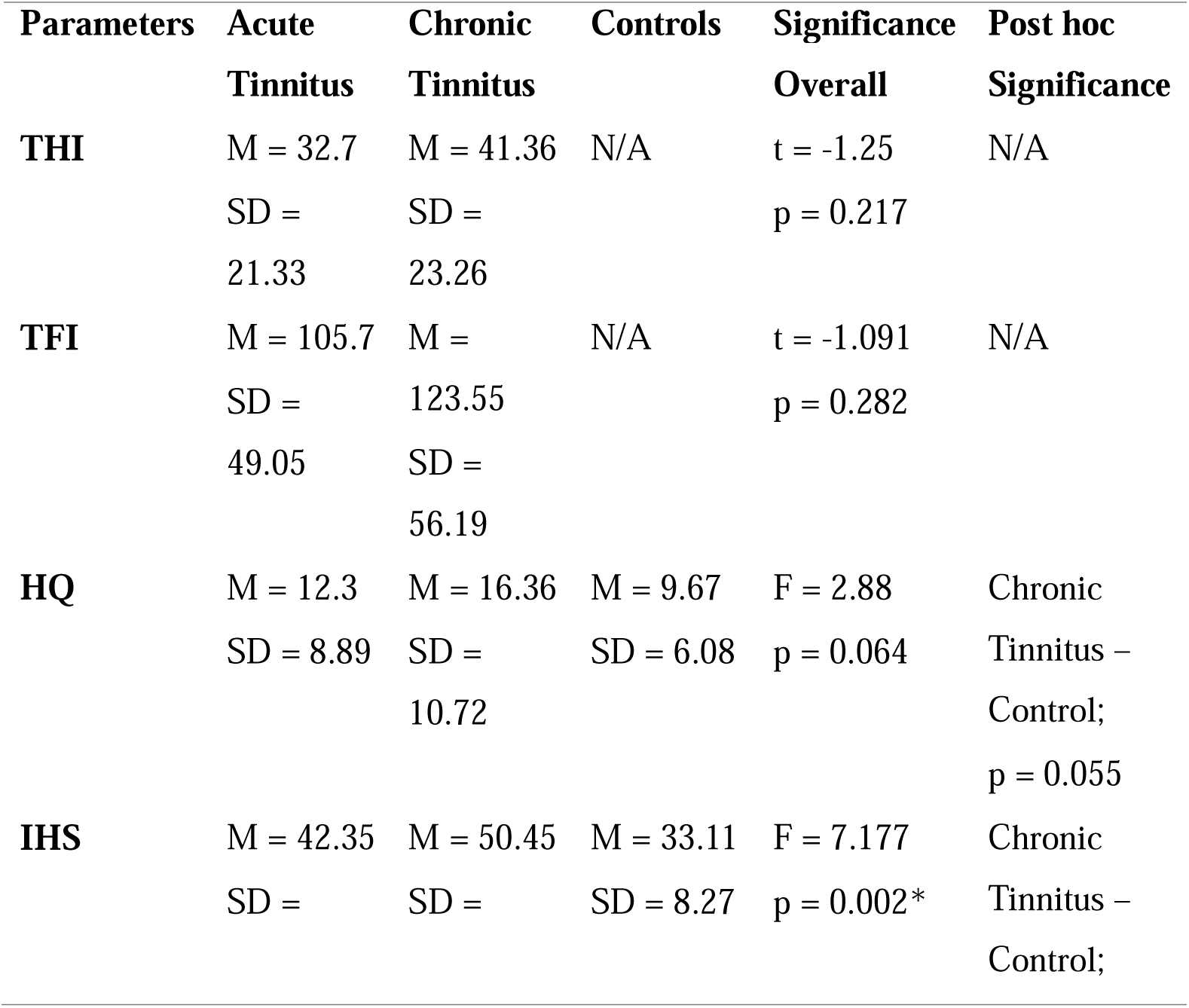
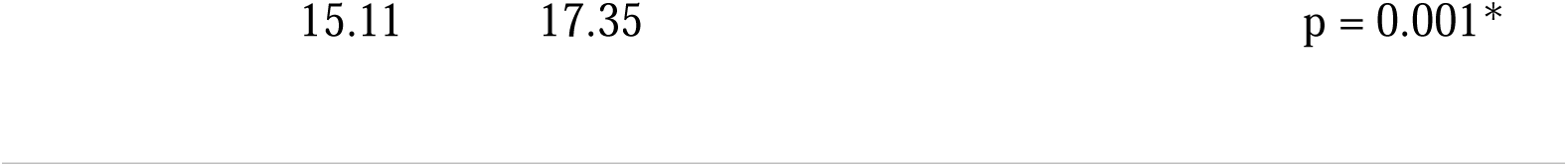
The participant’s symptom scores for both tinnitus and hyperacusis. THI - Tinnitus Handicap Inventory, TFI – Tinnitus Functional Index, HQ – Hyperacusis Questionnaire, IHS – Inventory of Hyperacusis Symptoms

| Parameters | Acute | Chronic | Controls | Significance | Post hoc |
| --- | --- | --- | --- | --- | --- |
|  | Tinnitus | Tinnitus |  | Overall | Significance |
| **THI** | M = 32.7<br>SD = 21.33 | M = 41.36<br>SD = 23.26 | N/A | $t = -1.25$<br>$p = 0.217$ | N/A |
| **TFI** | M = 105.7<br>SD = 49.05 | M = 123.55<br>SD = 56.19 | N/A | $t = -1.091$<br>$p = 0.282$ | N/A |
| **HQ** | M = 12.3<br>SD = 8.89 | M = 16.36<br>SD = 10.72 | M = 9.67<br>SD = 6.08 | $F = 2.88$<br>$p = 0.064$ | Chronic<br>Tinnitus –<br>Control;<br>$p = 0.055$ |
| **IHS** | M = 42.35<br>SD = | M = 50.45<br>SD = | M = 33.11<br>SD = 8.27 | $F = 7.177$<br>$p = 0.002^*$ | Chronic<br>Tinnitus –<br>Control; |

### P300 Amplitude changes between Acute, Chronic, and Control. Groups

For the relative P300 amplitude (defined by calculating the average amplitude across three conditions and then normalizing each condition by dividing its amplitude by this average to obtain the relative amplitude for each condition), no significant effect was seen for HFD F((2,62) = 0.184, p = 0.832) and NS (F(2,62) = 0.779, p = 0.463), but an effect was seen for LFD (F(2,62) = 3.121, p = 0.05) with a significant difference between Acute Tinnitus and Controls (p = 0.04) with Controls (M = 1, SD = 0.38) having a higher amplitude when compared to Acute Tinnitus (M = 0.75, SD = 0.33). Refer Figure 2 for further information of P300 amplitudes and Figure 3 for individual waveforms across groups and conditions.

**Figure 2.**
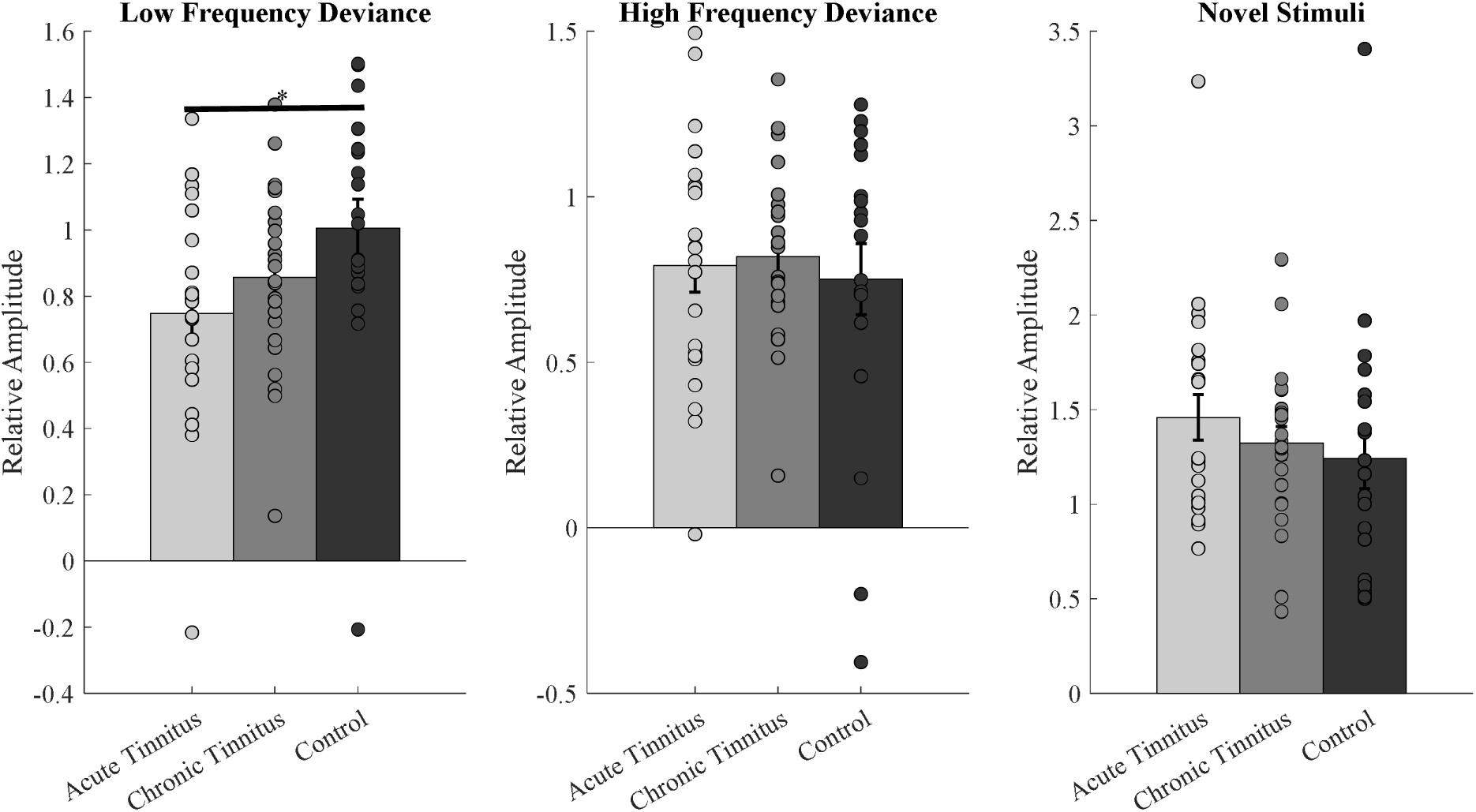
Relative P3 amplitude differences between Acute, Chronic, and Controls across conditions. The figures reveal the presence of significant difference between Acute Tinnitus and Controls for LFD and no significant differences across the groups for HFD and NS. The Asterix (*) indicates the presence of significant difference.

**Figure 3.**
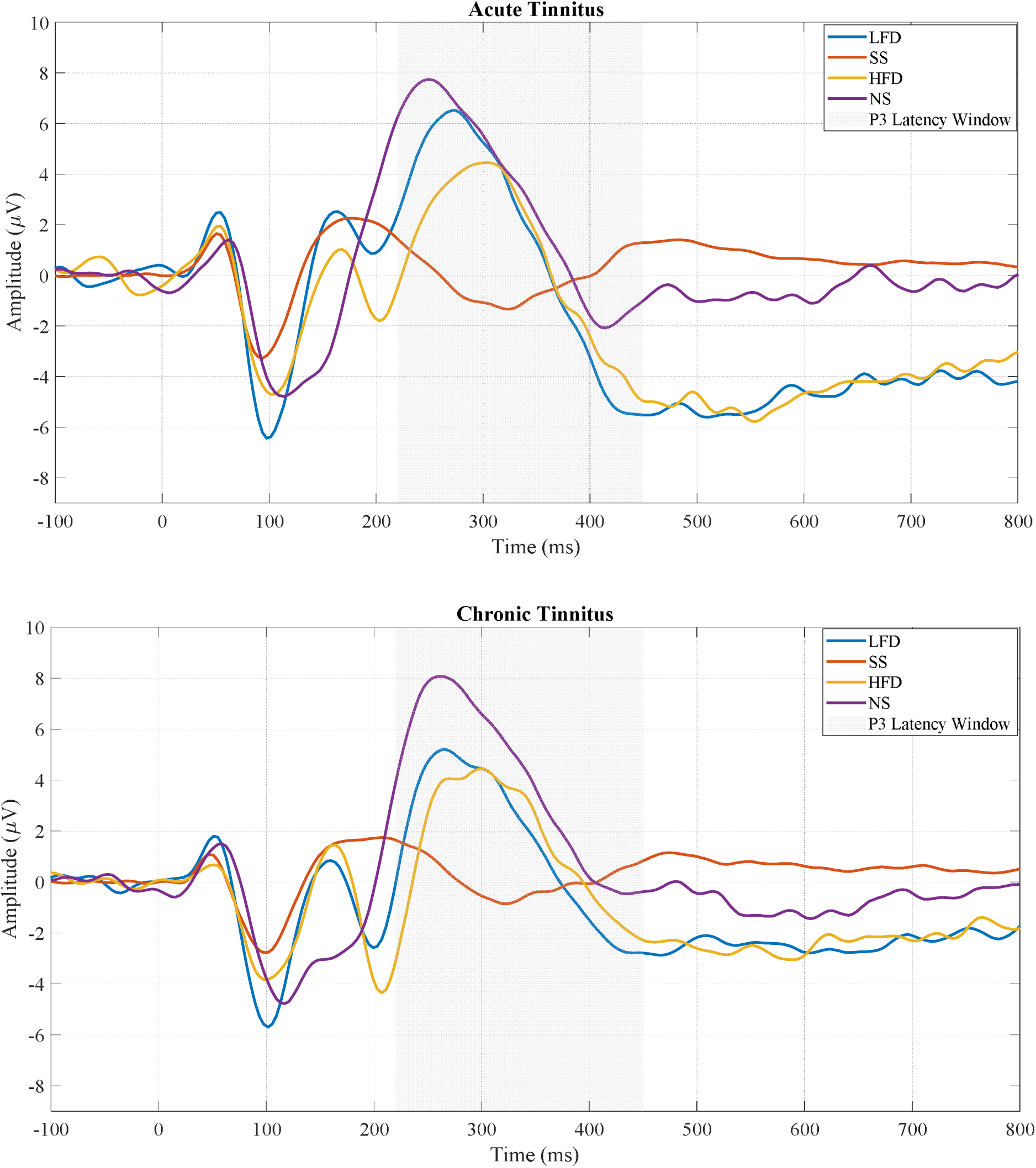
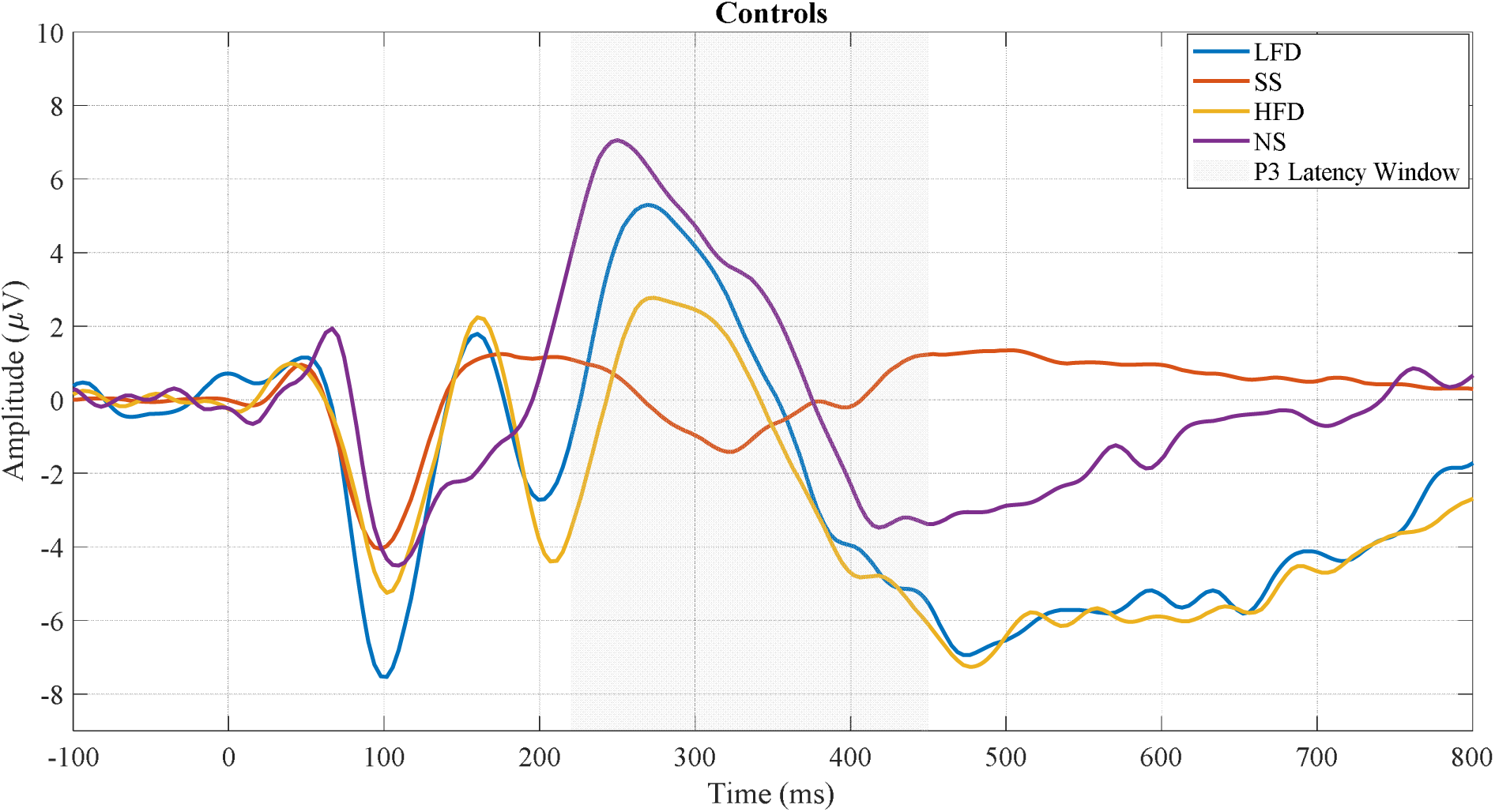
P300 waveforms between groups and across conditions. LFD: Low Frequency Deviance, SS: Standard Stimuli, HFD: High Frequency Deviance, NS: Novel Stimuli.

### P300 Amplitude changes between Acute Tinnitus and Post Acute Tinnitus

Upon paired longitudinal comparisons between the Acute and Post Acute group for the P300 amplitude, there was a non-significant trend for LFD (t(8) = 2.038, p = 0.076), a significant difference for HFD (t(8) = 3.47, p = 0.008), and a non-significant trend for NS (t(8) = 1.867, p = 0.099) with greater amplitude for Acute Tinnitus (LFD: M = 0.86, SD = 0.56, HFD: M = 0.73, SD = 0.38, novelty: M = 1.75, SD = 0.81) when compared to Post Acute Tinnitus (LFD: M = 0.62, SD = 0.4, HFD: M = 0.5, SD = 0.39, novelty: M = 1.54, SD = 0.8). Figure 4 illustrates the relative amplitude changes between Acute and Post Acute Tinnitus.

**Figure 4.**
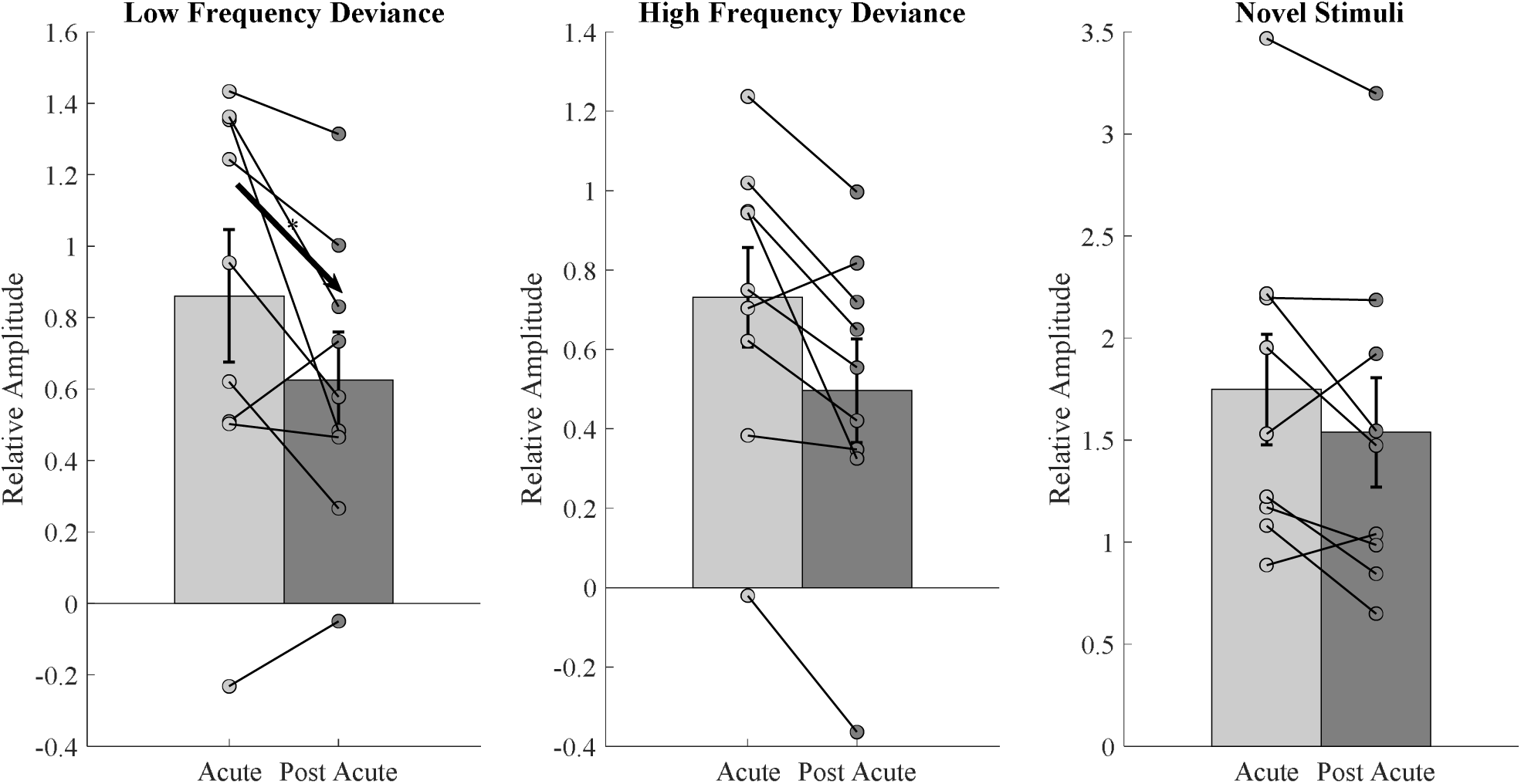
Relative P3 amplitude differences between Acute and Post Acute Tinnitus across conditions. Significant difference noted for LFD and non-significant trend noted for HFD and NS.

### Source reconstruction

An examination of source localisation was performed for both independent and matched groups. Currently, we are presenting solely the maximum or minimum t-statistics among the groups. As stated in the preceding section, we are exclusively reporting the source localisation activity for the groups that achieved statistical significance in the sensor space, specifically in the cross-sectional comparison between Acute Tinnitus and Controls for LFD, and the paired non-significant trend to significant differences between Acute and Post Acute Tinnitus for LFD, HFD, and NS. Between the Acute Tinnitus and Controls for LFD, the sources were localized in the inferior frontal gyrus. Concerning the longitudinal changes between Acute and Post Acute Tinnitus, the sources were localized in the anterior cingulate gyrus, inferior parietal lobe, and cingulate gyrus for HFD, LFD, and NS respectively. Figure 5 illustrates the source spaces for the above-mentioned group differences.

**Figure 5.**
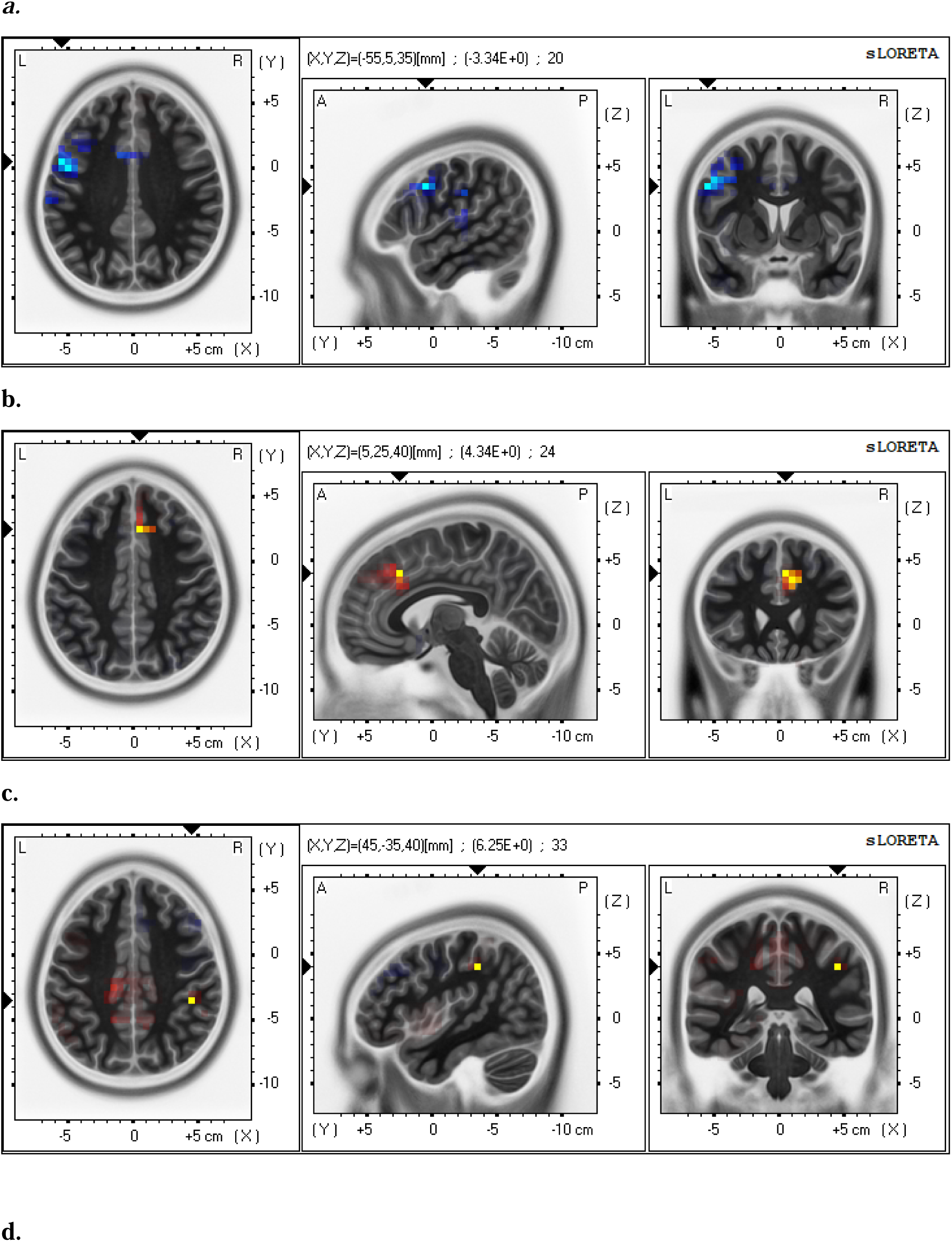
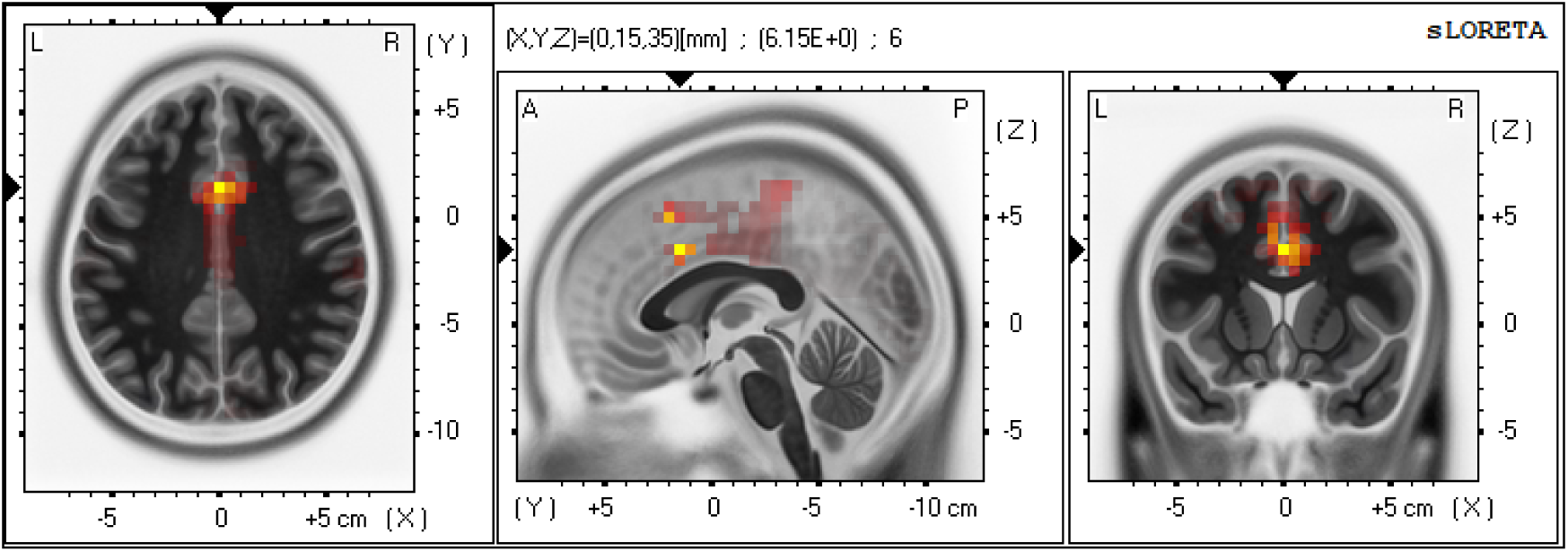
Source localization between group comparisons using sLORETA for peak P300 amplitude across conditions. The above illustrated sources indicate maximum t statistics. a. Source localized in the inferior frontal gyrus between Acute Tinnitus and Control for LFD. b. Source localized in the anterior cingulate cortex between Acute Tinnitus and Post Acute Tinnitus for HFD. c. Source localized in the inferior parietal lobe between Acute Tinnitus and Post Acute Tinnitus for LFD. d. Source localized in the anterior cingulate cortex between Acute Tinnitus and Post Acute Tinnitus for Novelty.

## Discussion

The primary findings of our data include alterations in P300 amplitude for LFD between Acute and Control as well as a general trend of decrease in P300 amplitude in Post Acute Tinnitus compared to Acute Tinnitus across the three conditions.

### Amplitude changes between Acute Tinnitus, Chronic Tinnitus, and Controls

Regarding P300 investigations in tinnitus, there is a paucity of literature in this domain, prompting us to draw inferences from the extensively researched Mismatch Negativity (MMN) studies related to tinnitus. The limited P300 studies, and more numerous MMN studies, in tinnitus have consistently shown reduced amplitudes of responses, at least to stimuli remote from the tinnitus frequency. With respect to the frequency deviants, the reduction in amplitude in Acute Tinnitus relative to Controls for LFD is concordant with previous P300 studies comparing Chronic Tinnitus with Controls at frequencies away from the tinnitus frequency. Gabr et al. (2022) conducted a study with the deviant stimuli being around 1050 Hz and the standard stimuli being around 1000Hz, they attributed the reduced P300 amplitude and prolonged latency to cognitive impairment and further implied attentional deficits in tinnitus due to possible involvement of the dorsolateral prefrontal cortex. A study by Shalaby et al. (2022) reported reduced P300 amplitude and prolonged latencies in individuals with tinnitus especially at a deviant frequency of 2 kHz. They attribute that the reason might be due to the impairment of central auditory processing and selective attention to the stimulus as a consequence of tinnitus. We further highlight MMN studies that have conducted deviance detection in relation to tinnitus due to the scarcity of literature related to P300 and tinnitus. Most of the MMN studies in tinnitus highlight the reduction of attention away from the tinnitus frequency when compared to at the tinnitus frequency (Yukhnovich et al., 2024).

In our study, we obtained a non-significant increase in P300 amplitude for Acute Tinnitus for novelty processing when compared to Controls, which we attribute to heightened salience (increased focussed attention) and distress associated with Acute Tinnitus during its initial phases. It is crucial to acknowledge that all existing implications stem from Chronic Tinnitus research, and to our knowledge, none have been documented in Acute Tinnitus. The sole MMN study in Acute Tinnitus by (Sedley et al., 2016) indicated that there were no significant differences between Acute and Chronic Tinnitus in intensity deviance detection, which is on par with our current findings from a cross-sectional comparison of both conditions, potentially attributing this to an absence of differences in attention allocation between the groups.

The source reconstruction revealed a reduction in the inferior frontal gyrus regions for Acute Tinnitus compared to Controls, for LFD. This has been attributed to overall reduction in distress when there is increased activity in the frontal regions which can be implicated with controls having reduced distress (Carpenter-Thompson et al., 2015). Furthermore, Lan et al. (2021) highlighted the presence of heightened attention towards the tinnitus in the acute stages of tinnitus when compared to the chronic stage which is due to the potential role of fronto-parietal attentional network. In our case, a similar argument can be considered possible for the Post Acute Tinnitus group as there is reduction of activity around the inferior parietal lobe in this group when compared to the Acute Tinnitus group for an incoming acoustic signal. This further implies that there is a high possibility of increased orientation of inferior parietal lobe activity to the tinnitus in the Post Acute Tinnitus group indicating the potential involvement of fronto-parietal attentional network during Post Acute Tinnitus (onset of chronification of tinnitus) facilitating its persistence. Another potential explanation associated with the function of the inferior frontal gyrus pertains to the prediction update mechanism. Todd & Robinson (2010) investigated the brain’s capacity to utilise contextual information for updating predictions and reducing prediction error. In an MMN study conducted by them, which sought to identify the differences in the deviance response between randomly generated deviance and linked deviance (non-randomized deviant stimulus generation), the results indicated a decrease in MMN amplitude to the linked deviance. This reduction may be attributed to successful anticipation of the linked deviance stimuli, thereby diminishing prediction errors and the MMN response. This anticipatory behaviour, according to them, is closely associated with the inferior frontal gyrus. Individuals with Acute Tinnitus may experience heightened attention and distress towards the tinnitus during its onset, which can diminish their focus on stimuli outside the tinnitus frequency. This reduction impairs the predictive update mechanism to LFD, leading to an altered amplitude in comparison to Controls.

### Post-Acute Tinnitus has a decreasing trend of P300 amplitude when compared to Acute Tinnitus

All three scenarios demonstrate either a reduced or trend in reduced P300 amplitudes in Post Acute Tinnitus relative to Acute Tinnitus, to all deviant types. The decrease in Post Acute Tinnitus relative to Acute Tinnitus corroborates our hypothesis that salience or novelty components are amplified during acute phases, subsequently decreasing and stabilising in the chronic stage. This pattern also aligns with frequency deviance, and our results emphasise the relatively intensified processing of novelty in contrast to the relatively reduced novelty and frequency deviance processing during the post-acute stages of tinnitus. The source reconstruction indicates a nearly significant elevation of activity in the Cingulate gyrus for both HFD and NS, as well as in the inferior parietal lobe for LFD, during Acute Tinnitus compared to the Post Acute Tinnitus groups.

### Novelty changes between Acute and Post Acute Tinnitus

Regarding novel stimulus responses, we interpret the current findings as an increased novelty-related activity in the Acute stage rather than a diminished activity in the Post Acute stage, given the direction of cross-sectional comparisons between Acute Tinnitus, Chronic Tinnitus, and Controls. The Controls tend to have a non-significant reduction of novelty when compared to both Acute Tinnitus and Chronic Tinnitus. Although the results were not statistically significant, we considered the direction of the amplitude changes across groups in the cross-sectional analysis to help interpret the patterns observed in the longitudinal comparisons. The aforementioned results regarding novelty may be attributed to heightened tinnitus-related distress during the acute phase, with the activity of cingulate gyrus strongly associated with distress. Vanneste et al. (2010) emphasised the potential role of the cingulate gyrus in tinnitus-related distress, given its crucial function within the limbic system.

Regarding the alleviation of tinnitus-related distress over time were corroborated by Vielsmeier et al. (2020), who emphasised that although the characteristics of tinnitus remained unchanged, there was a moderate decrease in tinnitus-related distress over time. Research conducted by Wallhäusser-Franke et al. (2017) revealed no alterations in distress and perceived loudness among persons with elevated distress at onset; however, a decrease in distress was noted in the low-level distress group, aligning with our present findings.

In addition to the cingulate cortex being linked to distress, it is also a crucial element of the attentional networks, which have been extensively related with tinnitus. Simonetti & Oiticica (2014) hypothesized increased activities in the regions of cingulate gyrus along with prefrontal cortex and insula which correlates with improved cognition regarding focused attention. The article emphasizes that individuals with tinnitus tend to show increased cingulate gyrus activity, indicating that their focused attention is predominantly directed towards tinnitus, similar to the attention tasks non-tinnitus individuals tend to focus on. A recent study by Vanneste et al. (2024) highlighted the presence of an on-and-off mechanism for individuals with intermittent tinnitus that is influenced by the pregenual anterior cingulate cortex. The onset of tinnitus is associated with increased theta activity and reduced functional connectivity with the auditory cortex, while the offset of tinnitus is facilitated by an increased alpha activity in the pregenual anterior cingulate cortex. Apart from these, the anterior cingulate cortex has also been associated with error detection or conflict monitoring as reported by Dali et al. (2022). The increased salience activity at the commencement of tinnitus can be ascribed to its novelty, as tinnitus serves as a novel stimulus that elicits a top-down response from the anterior cingulate cortex, functioning as a conflict detector. The heightened awareness of novel tinnitus during its onset correlates with an increased response to externally presented novel stimuli, as evidenced by a recent study on pain conducted by Hubbard et al. (2020a). This study identified an aberrant increase in the salience network in response to pain stimuli in fibromyalgia patients, suggesting a generalised hypervigilance to salient stimuli. This phenomenon is similarly observed in our Acute Tinnitus patients, who can either exhibit a generalised increase in hypervigilance towards any external salient/novel stimuli due to their heightened awareness or may even be the potential cause of the emergence of tinnitus.

The anterior cingulate cortex’s error detection property resembles the Bayesian predictive coding phenomenon, wherein Bayesian coding theory asserts that the brain probabilistically encodes sensory information through probability distributions. The brain constructs an internal model and juxtaposes it with external sensory input; greater similarity between the sensory input and the internal model results in reduced prediction error, while lesser similarity leads to increased prediction error (Knill & Pouget, 2004). The same has been applied to tinnitus research as well with predictive coding models formulated by De Ridder et al. (2014) and Sedley et al. (2016) where they state that the absence of auditory input from hearing loss prior to its onset creates a discrepancy between the predicted and the actual input, leading to a prediction error. To mitigate prediction error resulting from deafferentation, the brain often compensates for absent information, which may manifest as tinnitus. We wish to emphasise that the anterior cingulate cortex’s role in error detection for Acute Tinnitus may occur even during the precursor phase of tinnitus amidst hearing loss, which subsequently diminishes over time following the onset of tinnitus. These further associates the anterior cingulate cortex and novelty with the generation of tinnitus, aiming to minimise the prediction error between actual reduced input and expected input, in alignment with the model proposed by De Ridder et al. (2014). An alternative hypothesis suggests that the anterior cingulate cortex/salience may play a heightened role following the onset of tinnitus, wherein the tinnitus signal could induce an elevation in prediction error due to the discrepancy between anticipated sensory input (silence) and actual input (augmented spontaneous activity). This prediction error is interpreted as tinnitus, which diminishes over time as the internal model transitions from a default state of silence to increased spontaneous activity, as noted by Sedley et al. (2016). Our findings of reduced amplitude for novelty over a period of time suggests a decrease in the prediction error, accompanied by a decline in the activity of the anterior cingulate cortex during the same period further supports the model by Sedley et al. (2016) and in accordance with our principal hypothesis.

### LFD and HFD changes between Acute and Post Acute Tinnitus

The amplitudes for both LFD and HFD were showed either reduction or trend in reduction in the Post Acute stages of tinnitus. In contrast to the interpretation indicating possible adaptation to novelty in the post-acute phase, it has been demonstrated that P300 amplitudes for LFD were diminished in Acute Tinnitus compared to Controls cross sectionally which is in the opposite direction if we were to keep the cross-sectional comparison to indicate the direction of results. Thus, the observed direction diverges from that associated with novelty, necessitating the interpretation that both LFD and HFD exhibit reduced activity over time. Furthermore, the additional decrease in Post Acute Tinnitus relative to Acute Tinnitus may be ascribed to attention reallocations between frequencies. In this instance, contrary to other studies, we are not directly linking reduced cognitive performance or attention deficiencies in patients with tinnitus Gabr et al. (2022) but rather suggesting a pre-conscious diversion of attention from external stimuli due to an internal concentration on the tinnitus. The inferior parietal cortex that was visualized in the source space for LFD with reduced inferior parietal activity over time is associated with the fronto-parietal attentional network, where recent research emphasises the heightened focused attention on tinnitus, hence increasing the fronto-parietal attentional network towards the tinnitus and thereby reducing attention towards external stimuli (Hubbard et al., 2020b).

### Reduced attention to external stimuli may facilitate the persistence of tinnitus

In the current research, we concentrated solely on the cortical region with the greatest t statistic as our region of interest, potentially restricting the evaluation of other brain regions that may demonstrate independent activity or possess pertinent functional connections.

However, evaluating the current investigation at face value, it is intriguing to observe the comparisons between Acute and Chronic Tinnitus both cross-sectionally and longitudinally, the longitudinal comparisons yielded significantly more robust results, demonstrating non-significant trend to significant reductions in relative amplitude across conditions from Acute to Post Acute stages. The inferior parietal lobe which was the other region of interest in this study apart from the anterior cingulate cortex has been known to contribute to both novelty processing (Kiehl et al., 2001) and in P3b generations too (Volpe et al.,2007). The role of the inferior parietal cortex can be interpreted in two manners: either as a heightened awareness of the tinnitus at its inception or as a reduced attentiveness to external stimuli throughout the chronic stages of tinnitus. We would favour the latter based on the evidence from the current study. In addition to the alterations in the inferior parietal lobe, we observed a decrease in P300 amplitudes in both target deviant circumstances at sensor space, indicating attention relocation between frequencies (Volpe et al., 2007).

It is imperative to note that the increased activity in the salient processing of the tinnitus signal increases attention orientation to any externally incoming salience/novel stimulus. But it is the contrast for frequency deviance detection with reduced activity in the inferior parietal lobe during the later stages or Post Acute stages of tinnitus, thereby reducing the attention to the external stimuli, due to the increase in attention towards the tinnitus. We further propose that there is a possibility of the anterior cingulate cortex to play a role in tinnitus formation through error detection in the *precursor stages*, leading to tinnitus as a mechanism to mitigate prediction error. The other possibility is that the prominence or novelty of tinnitus is affected by the anterior cingulate cortex, which functions as an error detector at the onset of tinnitus, subsequently engaging additional attentional networks that reduce focus on external stimuli, thereby amplifying concentration on tinnitus and ensuring its persistence. These findings align with the initial resting state EEG results, which emphasise the anterior cingulate cortex for cross-sectional differences between Acute and Chronic Tinnitus, and the inferior parietal cortex for longitudinal differences between Acute and Post-Acute Tinnitus, further underscoring the collaborative role of the anterior cingulate cortex and inferior parietal cortex in the onset and persistence of tinnitus.

Figure 6 depicts a newly proposed account of tinnitus persistence based on the attentional orientation towards the tinnitus further explaining its onset and persistence. A significant drawback of this study is the Chronic Tinnitus cohort, which could not be matched for high-frequency hearing, hence preventing the current study from interpreting the cross-sectional comparisons of Chronic Tinnitus with the other two groups.

**Figure 6.**
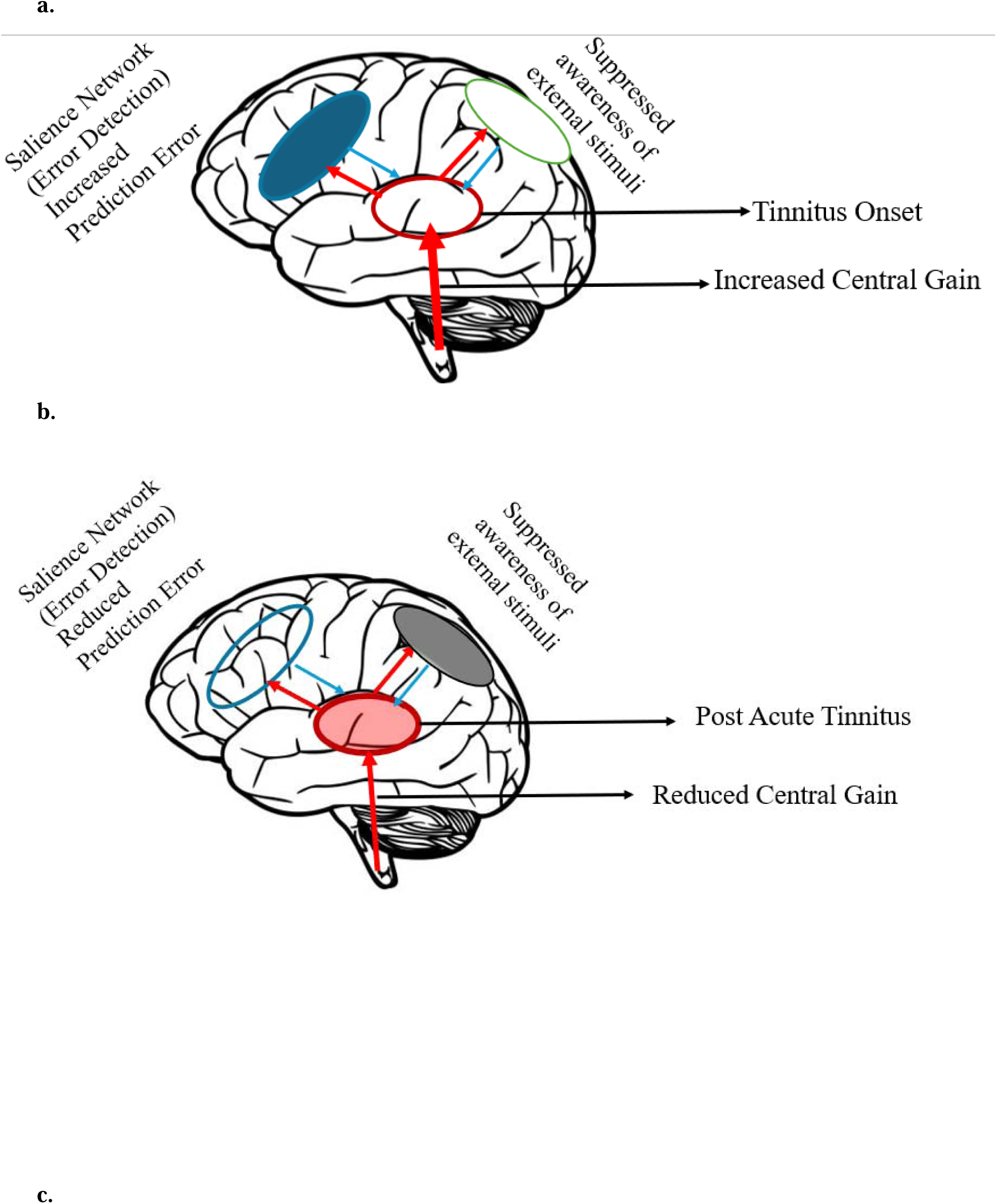
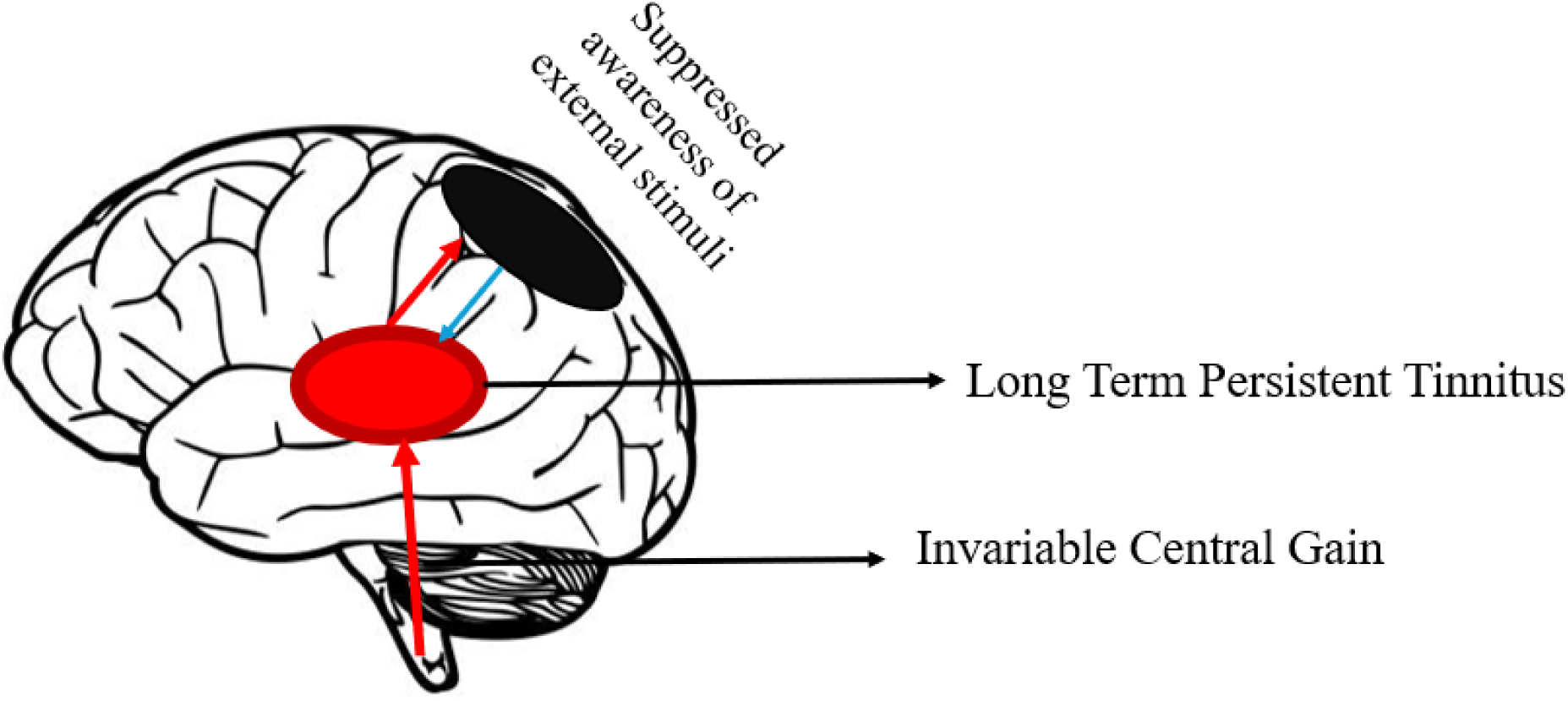
Illustration of tinnitus mechanism based on stimulus novelty. The three figures illustrate changes in tinnitus mechanism post onset of tinnitus. There are three major elements included in the figure which include anterior cingulate cortex (blue) highlighting error detection, inferior parietal lobe (black) highlighting increased attention orientation towards the tinnitus, auditory cortex (red) highlighting the tinnitus activity, and increased central gain (red arrow) which increases during the onset and regresses to the mean during the Post Acute and Chronic stages. *a.* Acute Tinnitus: There is increased activation of salience network around the time of the onset of tinnitus which either may cause tinnitus or be a consequence of tinnitus. There is increased prediction error around this time concurrently activating the attentional networks which includes the inferior parietal cortex that starts to reorganize, diminishing focus on external stimuli and redirecting it towards the tinnitus. The central gain increases during this stage that results in tinnitus onset. *b.* Post Acute Tinnitus: The propagation of tinnitus diminishes prediction error by altering the mental model, shifting the default prediction from quiet to tinnitus, so causing the salience network to reduce the error detection that was present around the onset of tinnitus. Conversely, the attentional networks (through fronto-parietal attentional networks) enhance the focus on the tinnitus while diminishing the perception of exterior stimuli. Central gain reduces as a function of regression to the mean. *c.* Chronic Tinnitus: The mental model has transitioned from default silence to tinnitus, thereby minimizing prediction error. Concurrently, the attentional networks (specifically sustained attention) experience a permanent alteration in connectivity, which enhances focus on tinnitus while diminishing attention to external stimuli, thus facilitating a persistent perception of tinnitus. Central gain remains constant indicating the tinnitus activity remaining constant over time after initial reduction.

### Conclusion

Previous research has proposed potential processes for the generation and persistence of tinnitus, either independently or through cross-sectional comparisons between acute and chronic tinnitus. The primary areas of focus have been the prefrontal, parietal, and insular regions in the generation and maintenance of tinnitus. We opted to examine our hypothesis using a novelty based P300 paradigm, which is typically elicited from both prefrontal and parietal regions, to investigate the distinction between acute and chronic tinnitus and to corroborate earlier findings. This research likely represents the first study employing a longitudinal design that includes a follow-up on Acute Tinnitus six months post-onset of Chronic Tinnitus. Our data indicate a significant drop in P300 amplitude during the chronic stage, highlighting the substantial influence of the anterior cingulate cortex/salience network in possible generation of tinnitus and inferior parietal lobe in the persistence of tinnitus.

